# Human Mitochondrial AAA+ ATPase SKD3/CLPB forms Nucleotide-Stabilized Dodecamers

**DOI:** 10.1101/2021.12.16.472980

**Authors:** Zachary Spaulding, Indhujah Thevarajan, Lynn G. Schrag, Lejla Zubcevic, Anna Zolkiewska, Michal Zolkiewski

## Abstract

SKD3, also known as human CLPB, belongs to the AAA+ family of <u>A</u>TPases <u>a</u>ssociated with various <u>a</u>ctivities. Mutations in the *SKD3/CLPB* gene cause 3-methylglutaconic aciduria type VII and congenital neutropenia. SKD3 is upregulated in acute myeloid leukemia, where it contributes to anti-cancer drug resistance. SKD3 resides in the mitochondrial intermembrane space, where it forms ATP-dependent high-molecular weight complexes, but its biological function and mechanistic links to the clinical phenotypes are currently unknown. Using sedimentation equilibrium and dynamic light scattering, we show that SKD3 is monomeric at low protein concentration in the absence of nucleotides, but it forms oligomers at higher protein concentration or in the presence of adenine nucleotides. The apparent molecular weight of the nucleotide-bound SKD3 is consistent with self-association of 12 monomers. Image-class analysis and averaging from negative-stain electron microscopy (EM) of SKD3 in the ATP-bound state visualized cylinder-shaped particles with an open central channel along the cylinder axis. The dimensions of the EM-visualized particle suggest that the SKD3 dodecamer is formed by association of two hexameric rings. While hexameric structure has been often observed among AAA+ ATPases, a double-hexamer sandwich found for SKD3 appears uncommon within this protein family. A functional significance of the non-canonical structure of SKD3 remains to be determined.

---

Suppressor of potassium transport defect 3, SKD3, also known as caseinolytic peptidase B homolog CLPB, belongs to the AAA+ family of <u>A</u>TPases <u>a</u>ssociated with various <u>a</u>ctivities.^1,2^ The function of SKD3/CLPB is not fully understood and its relation to the well-studied protein disaggregases ClpB/Hsp104 has been under debate. While expression of the disaggregases ClpB/Hsp104 is limited to microorganisms and plants^3^, SKD3 is only found in vertebrates.^4^ SKD3 contains the N-terminal mitochondrial targeting sequence, followed by an ankyrin-repeat domain and a single ATP-binding AAA+ module.^4^ In contrast, members of the ClpB/Hsp104 family contain two AAA+ modules separated by a characteristic coiled-coil domain, which couples the disaggregation function with aggregate recognition by the co-chaperones DnaK/Hsp70.^5,6^ The coiled-coil domain is missing in SKD3. While ClpB/Hsp104 are primarily cytosolic and their isoforms are located in the mitochondrial matrix, SKD3/CLPB localizes to the mitochondrial intermembrane space (IMS).^7^ The mature form of SKD3/CLPB is produced upon its import into the IMS through proteolytic processing of the N-terminal targeting sequence.^8^

Hereditary mutations in the human *SKD3/CLPB* gene have been linked to various pathologies with a broad phenotypic spectrum, including 3-methylglutaconic aciduria, congenital neutropenia, cataract, intellectual disability/developmental delay, progressive brain atrophy, and movement disorder.^9-13^ Recently, *de novo SKD3/CLPB* mutations have been also associated with congenital neutropenia.^14,15^ Notably, the *SKD3/CLPB* expression is elevated in acute myeloid leukemia (AML) and further upregulated upon acquisition of resistance to apoptosis-inducing treatment.^16^ In AML cells, SKD3/CLPB is involved in maintaining the mitochondrial cristae structure and SKD3 ablation promotes sensitivity to pro-apoptotic drugs.^16^ Since SKD3 reactivates selected aggregated proteins *in vitro*, it was postulated to function as a molecular chaperone/disaggregase in the IMS.^4^

We have previously shown that SKD3 forms ATP-dependent high-molecular weight complexes in the IMS.^7^ The composition and structure of the SKD3-containing molecular assemblies in the IMS are unknown. To begin closing that gap in knowledge, we produced a recombinant mature form of human SKD3 in *Escherichia coli*. We used dynamic light scattering, sedimentation equilibrium, sedimentation velocity, and negative-stain electron microscopy to obtain the first structural information on the oligomeric assembly of human SKD3.

Purified recombinant SKD3 corresponding to the mature IMS form (residues 127-707, monomer molecular weight 65,810 Da) displayed a robust ATPase activity of 27 ± 3 nmol ATP *per* nmol SKD3 *per* min, which agrees with the previously reported activity^4^ and is several-fold higher than the ATPase activity of *E. coli* ClpB^17^. The SKD3 ATPase did not respond to 0.1 mg/ml α-casein, a well-known pseudo-substrate and activator of ClpB/Hsp104, again in agreement with the previous results.^9^ The lack of casein-induced activation the SKD3 ATPase suggests that its substrate-recognition mechanism may differ from that of the disaggregases ClpB/Hsp104.

The particle-size distributions determined with dynamic light scattering (DLS) showed that the apparent hydrodynamic (Stokes) diameter of SKD3 particles did not exceed 1 nm at protein concentration below 0.1 mg/ml (Fig. 1A). However, the apparent SKD3 size increased to over 10 nm in diameter at a higher protein concentration (Fig. 1B) or in the presence of nucleotides: ATP, ADP, or the non-hydrolysable ATP analogue adenylyl-imidodiphosphate (AMP-PNP) (Fig. 1C, D, E). We conclude that SKD3 undergoes self-association into oligomers, which are stabilized by adenine nucleotides. The SKD3 oligomer stabilization does not require ATP hydrolysis and is induced by the diphosphate as well as the triphosphate nucleosides.

**Figure 1.**
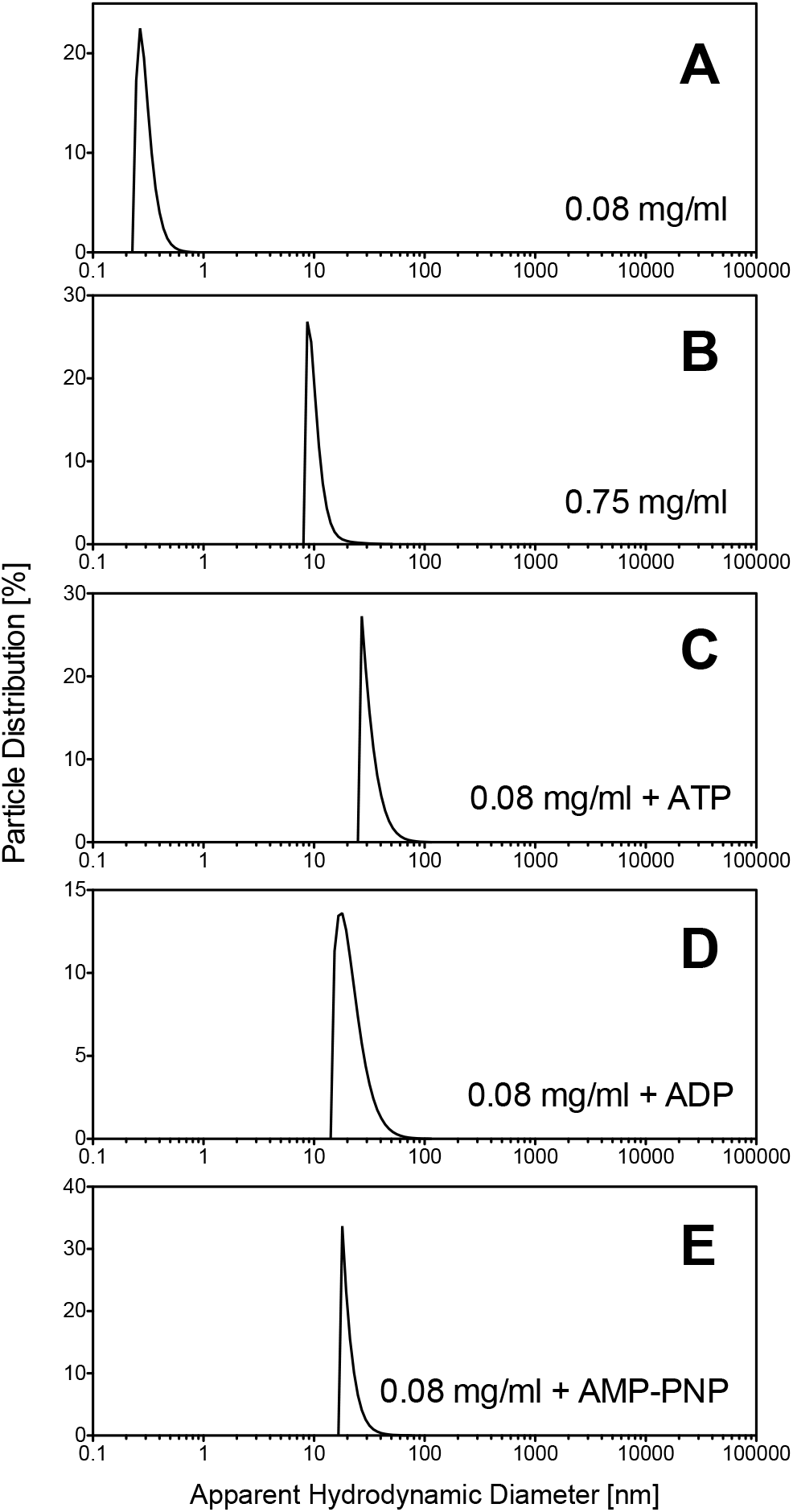
Dynamic light scattering analysis of SKD3/CLPB samples. The SKD3 particle-size distributions were measured at the indicated protein concentration without nucleotides or with 1.7 mM ATP, ADP, or AMP-PNP. Shown are representative results from 2 repeated experiments.

To determine the molecular weight of the SKD3 species detected with DLS, we performed sedimentation equilibrium experiments at low and high protein concentration, in the absence and presence of ADP. As shown in Fig. 2A, the protein concentration gradient for 0.08 mg/ml SKD3 in the absence of ADP was consistent with monomeric SKD3. At 1.2 mg/ml SKD3 in the presence of 1 mM ADP, the protein molecular weight increased approximately 12-fold (Fig. 2B). The protein concentration gradient for SKD3 at 1.2 mg/ml without a nucleotide could not be approximated with a single-species model (data not shown), which suggests that the oligomers formed at high protein concentration in the absence of nucleotides are less stable than those produced in their presence. Importantly, the results in Fig. 2B show, for the first time, that SKD3 forms dodecameric assemblies.

**Figure 2.**
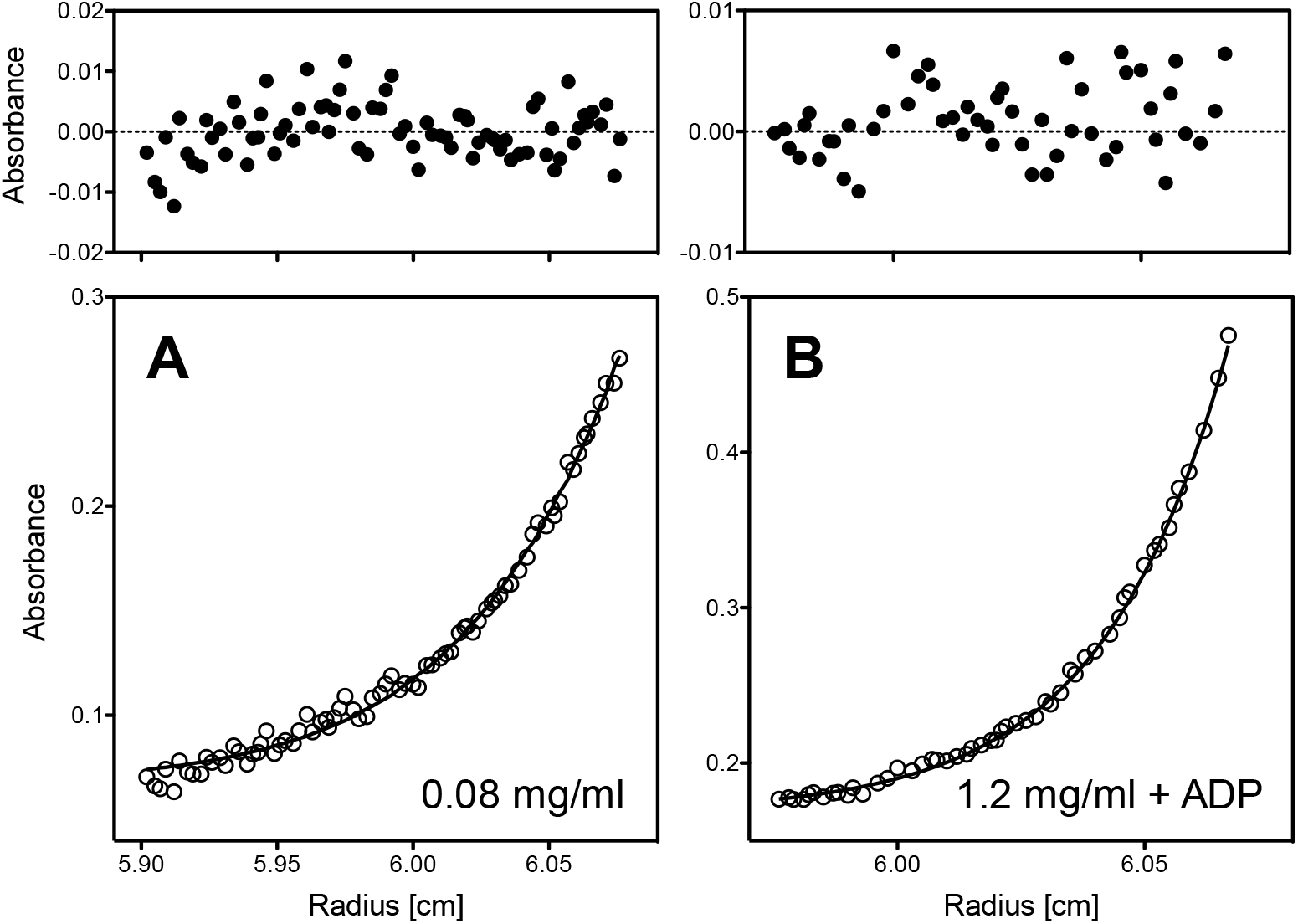
Sedimentation equilibrium analysis of SKD3/CLPB. Bottom panels: shown are the protein concentration gradients (open circles) and the fits corresponding to single-species model (solid lines). Top panels: shown are the residuals of the fits (A_exp_ – A_model_). The concentration gradients were measured at 4 ºC after the sedimentation equilibrium was reached at 20,000 rpm with 0.08 mg/ml SKD3 (**A**) or at 8,000 rpm with 1.2 mg/ml SKD3 and 1 mM ADP (**B**). The protein absorption was measured at 235 nm (**A**) or 295 nm (**B**). The single species fits gave an apparent molecular weight of 65,780 Da with the non-ideality coefficient of 1.4·10^−5^ (**A**) and 788,230 Da with the non-ideality coefficient -6·10^−9^ (**B**).

To obtain further insight into the structure of the SKD3 dodecamer, we performed negative-stain electron-microscopy imaging of SKD3 in the presence of 1 mM ATP. The sample images showed uniformly distributed large particles, which was consistent with the stabilizing effect of the nucleotide (Fig. 3, top panel). Image-class analysis and averaging visualized cylinder-shaped particles with an open central channel along the cylinder axis (Fig. 3, bottom panels). The SKD3 particle top view showed a ring structure with the diameter of ∼13 nm and a central pore with the diameter of ∼2 nm. The ring dimensions determined for SKD3 agree with those found for the hexameric assembly of bacterial ClpB^18,19^ and other ring-forming hexameric AAA+ ATPases^20^. Interestingly, the SKD3 side view showed a ∼15 nm-tall and apparently triple-tiered structure with a top-bottom symmetry (Fig. 3, bottom panels).

**Figure 3.**
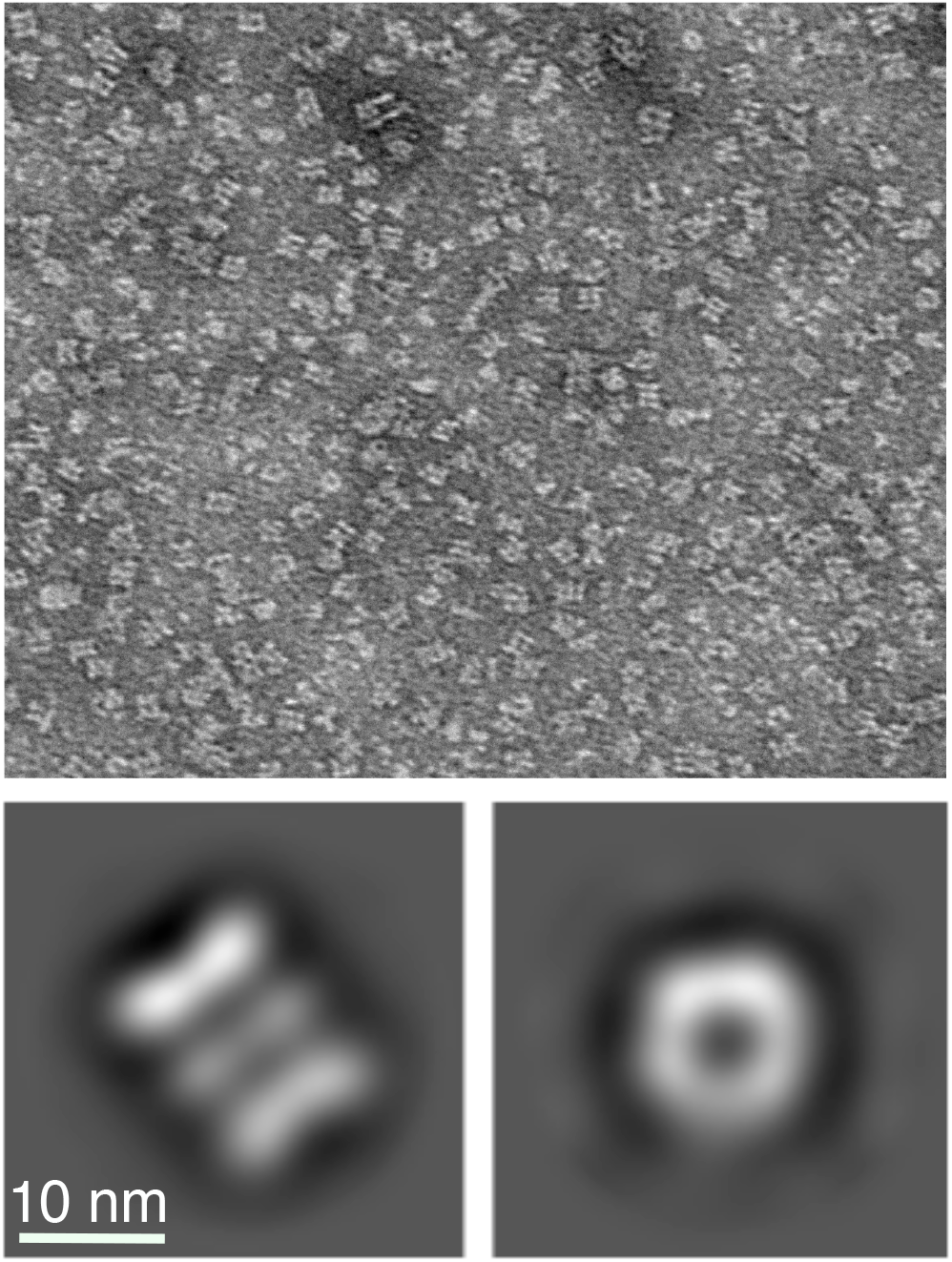
Negative-stain electron microscopy images of SKD3/CLPB. The top panel shows a raw electron micrograph of negatively stained 0.6 mg/ml SKD3 with 1 mM ATP at 30,000-fold magnification. The bottom panels show the side view (left) and top view (right) of the image-class analysis and averaging from the total of 12,548 particles.

The SKD3 oligomer molecular weight as determined by sedimentation equilibrium (Fig. 2B) indicates that the particle visualized in Fig. 3 contains twelve SKD3 subunits. Given the particle diameter, which approximates the diameter of a canonical hexameric AAA+ ring, the twelve SKD3 subunits are likely arranged in two ring-shaped hexamers stacked against each other. The side-view symmetry suggests that the association of two SKD3 hexamers occurs either “head-to-head” through the N-terminal ankyrin-repeat domains or “tail-to-tail” through the C-terminal AAA+ domains. Furthermore, in previous imaging analyses of hexameric AAA+ ATPases, the number of side-view tiers corresponded to the number of distinct ring structures formed by individual protein domains.^18,21-23^ Thus, a double-hexamer sandwich of a two-domain SKD3 might appear as a four-tier particle. Unexpectedly, in the SKD3 dodecamer side view, we observed three distinct structural rings instead of four, which suggests that the ring stacking might involve an overlap or intertwining of the SKD3 domains from the opposite rings. The detailed SKD3 domain arrangement in the dodecameric complex will need to be refined with future structural studies at higher resolution.

To further characterize the SKD3 dodecamers, we performed sedimentation velocity experiments for SKD3 with 1 mM ATP or without nucleotides. The maximum of the apparent sedimentation coefficient distribution curve in Fig. 4 defines the sedimentation coefficient of the predominant molecular species in solution.^24^ In the presence of ATP, SKD3 sedimented as a 16.2 S particle (Fig. 4A). The predicted sedimentation coefficient for a solid spherical particle with the molecular weight of the SKD3 dodecamer (792 kDa) is 27.9 S, which is significantly higher than the observed value. A reduction of the sedimentation rate could arise from a significant deviation from a spherical shape. However, the height/diameter ratio for the particles visualized in Fig. 3 is ∼1.15, indicating that the shape of an SKD3 dodecamer does not strongly deviate from a sphere. Alternatively, sedimentation rate may be strongly reduced if protein complexes form hollow particles with internal solvent-filled voids. Such hollow particles produce anomalously high frictional forces, which oppose the force of centrifugation. Indeed, a top view of the SKD3 dodecamer reveals a hollow channel/cavity (Fig. 3). Thus, a strong reduction of the sedimentation coefficient for the SKD3 dodecamers, as compared to that of a solid sphere with the same molecular weight, could be explained by the voids in their molecular structure.

**Figure 4.**
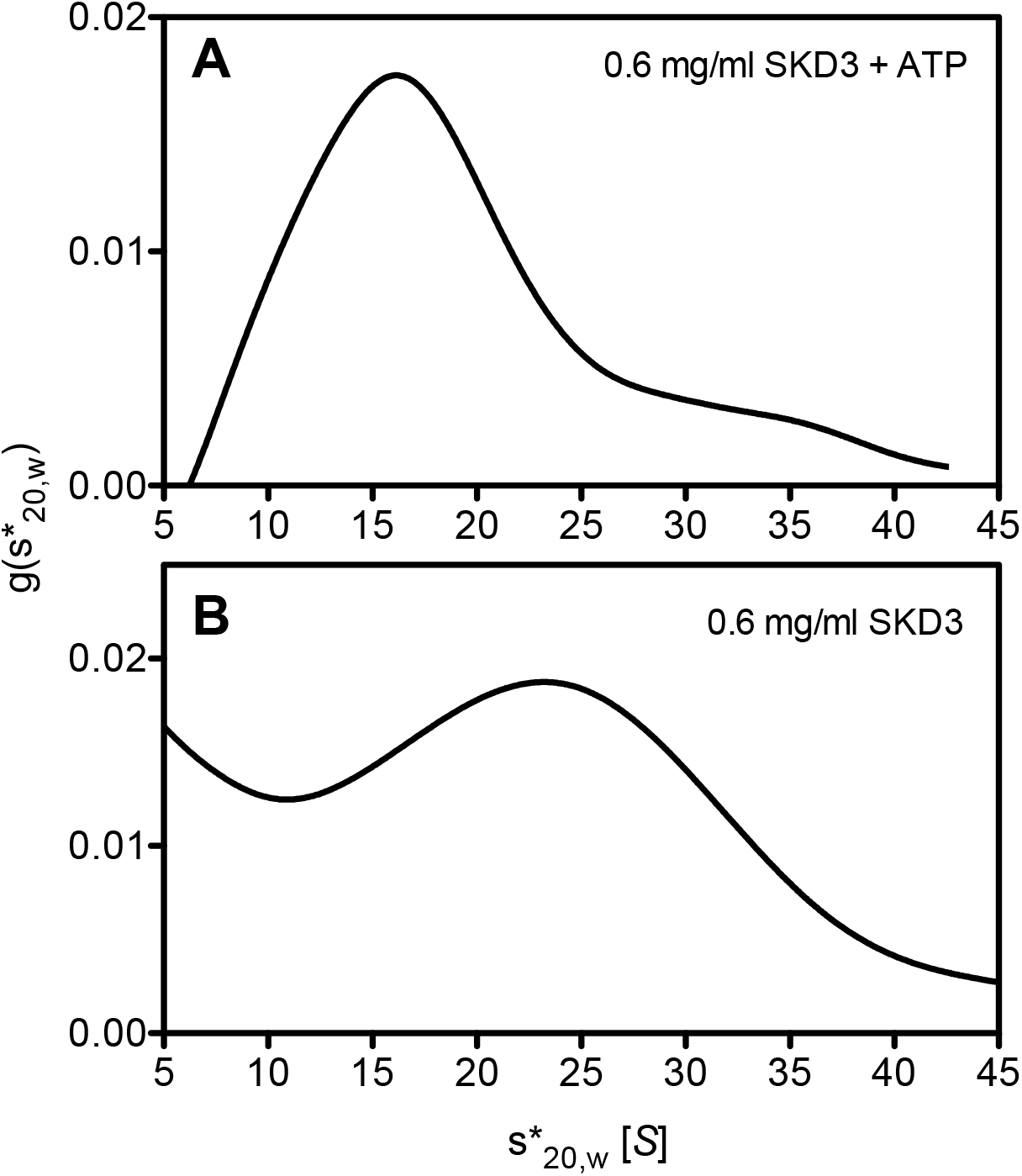
Sedimentation velocity analysis of SKD3/CLPB. Shown are the apparent sedimentation coefficient distributions g(s^*^_20,w_) vs. the sedimentation coefficient s^*^_20,w_ in Svedberg units (*S*) obtained from the time-derivative analysis^24^. The sedimentation experiments were performed at 20 ºC and 35,000 rpm. The progress of SKD3 sedimentation at 0.6 mg/ml was monitored with the interference optics in the presence of 1 mM ATP (**A**) or with absorbance at 280 nm in the absence of the nucleotide (**B**).

The sedimentation coefficient distribution for nucleotide-free SKD3 (Fig. 4B) differs significantly from that found in the presence of ATP (Fig. 4A). The distribution is broadened, and its maximum indicates that the predominant species sediments as a 23.2 S particle, which is significantly higher than the sedimentation coefficient of 16.2 S in the ATP-bound state. This result demonstrates a significant conformational change in the SKD3 oligomers induced by ATP. The sedimentation rate of SKD3 oligomers formed in the absence of nucleotides better approximates the sedimentation of solid spherical particles than the rate observed in the presence of ATP. It is also possible that the SKD3 oligomers formed in the absence of nucleotides are heavier, i.e., contain a higher number of subunits, as was observed previously for the bacterial ClpB.^19^

In this study, we obtained the first structural information for human SKD3/CLPB. We discovered that SKD3 forms nucleotide-stabilized dodecamers and provided the first EM image analysis for the dodecamers in the most physiologically relevant ATP-bound state. While hexameric structures have been observed often among other AAA+ ATPases^2,25^, the apparent double-hexamer sandwich found for SKD3 is uncommon within this protein family. A functional significance of the non-canonical structure of SKD3 remains to be determined.

In spite of its non-canonical double-hexamer assembly, the dodecameric SKD3 contains a characteristic structural feature that is essential for biological activity of AAA+ ATPases: an open central pore/channel (see Fig. 3). Among the variety of AAA+ functions, an insertion or translocation of substrates inside the central channel appears to be a common mechanistic principle of this diverse protein family.^2,25,26^ An identity of IMS-localized macromolecules that are processed in the SKD3 channel, a mechanism of SKD3-substrate interaction, and biological consequences of SKD3-driven substrate processing remain to be discovered.

## Materials and methods

### Protein expression and purification

The human SKD3/CLPB cDNA fragment corresponding to the residues 127-707 was PCR amplified to include *BamH*I and *Not*I restriction sites at the 5’ and 3’ ends, respectively, using the RC203013 vector (OriGene Technologies) as the template. The PCR product was digested with *BamH*I and *Not*I, and subcloned into the pOP3MT vector (plasmid #112601, Addgene). Recombinant mature SKD3 was produced as a TEV-cleavable fusion with the N-terminal Avi-His8 and maltose binding protein (MBP) tags using a procedure similar to that of Cupo and Shorter^27^, with modifications described below.

The pOP3MT plasmid containing the sequence of the mature form of SKD3 was transformed into *Escherichia coli* BL21(DE3) (Millipore-Sigma). Cells were cultured in 2 × 5 ml LB broth with 100 μ g/ml ampicillin for 16 hours at 30 °C with shaking, then transferred to 1 L of LB with ampicillin in a 4-L baffled Erlenmeyer flask and cultured with shaking at 37 °C for an additional 2.5 h. Induction was performed by supplementing the media with 1 mM IPTG and culturing the cells at 15 °C for 16 hours with shaking. Cells were collected by centrifugation at 4,500 × g for 30 min at 4 °C and then disrupted by sonication in lysis buffer (40 mM HEPES-KOH pH 7.4, 500 mM KCl, 20% (w/v) glycerol, 10 mM MgCl_2_, 2 mM β-mercaptoethanol, and a protease inhibitor cocktail (Sigma) in the amount 1 ml/100 ml buffer). Lysates were cleared via centrifugation at 30,000 × g at 4 °C for 20 min. Nucleic acids were precipitated from cleared lysates with 0.04% polyethyleneimine and pelleted by centrifugation at 20,000 × g for 25 min at 4 °C. The supernatant containing SKD3-MBP fusion protein was incubated with 4 ml amylose resin column overnight at 4 °C. The column was washed with 10 bed volumes of wash buffer (40 mM HEPES-KOH pH 7.4, 500 mM KCl, 10% (w/v) glycerol, 10 mM MgCl_2_, 2 mM β-mercaptoethanol) at 4 °C. Protein was eluted with amylose elution buffer (50 mM Tris-HCl pH 8.0, 300 mM KCl, 10% (w/v) glycerol, 10 mM MgCl_2_, 2 mM β-mercaptoethanol, 10 mM maltose.) SKD3-MBP fusion protein was cleaved with a 6His-tagged AcTEV™ protease (Thermo Fisher) at 4 °C for 96 h in the amylose elution buffer. After the cleavage, the sample was supplemented with 10 mM imidazole and incubated with 1 ml Ni-NTA resin (Qiagen) for 1 hour at 4 °C. In SDS-PAGE analysis, cleaved SKD3 was found in the wash fractions obtained with 20 mM sodium phosphate buffer, pH 7.4, 300 mM NaCl, 25 mM imidazole, 2 mM β-mercaptoethanol. The SKD3-containing fractions were collected and dialyzed into buffer A (50 mM HEPES-KOH pH 7.5, 200 mM KCl, 10 mM MgCl_2_, 1 mM EDTA, 2 mM β-mercaptoethanol). Protein concentration was determined by measuring absorbance at 280 nm, using the absorption coefficient 0.796 cm^2^/mg (calculated with Protparam, https://web.expasy.org/protparam/).

### Dynamic light scattering

DLS experiments were performed at room temperature with an Anton Paar Litesizer 500 instrument equipped with a Univette 50-μl sample compartment. SKD3 stock solution was diluted in buffer A to the desired concentration. For the experiments with nucleotides, the samples were incubated for 10 min at room temperature after adding the nucleotide. The final particle-size distributions were obtained from averaging of 25 correlation-function decays.

### Analytical ultracentrifugation

Sedimentation equilibrium and sedimentation velocity experiments were performed with a Beckman Optima XL-I analytical ultracentrifuge equipped with the interference and absorption detection systems and a four-position AN-Ti rotor. For sedimentation equilibrium experiments, 100 μl of SKD3 solution and 120 μl of buffer A were loaded to the right and left sectors, respectively, of the 6-channel analytical cell. The samples were equilibrated at 4 ºC at the speed indicated in Figure 2. To monitor the progress towards equilibrium, radial scans were performed every 6 hours. The final data collection was performed after 50-60 h. The data analysis was performed using the single-species model built into the Origin software supplied with the instrument. Partial specific volume of SKD3 (0.737 ml/g) and the buffer density were calculated using Sednterp software by John Philo (http://www.jphilo.mailway.com). For sedimentation velocity experiments, 300 μl of SKD3 solution and 320 μl of buffer A were loaded to the right and left sector, respectively, of the double-sector analytical cell. After equilibration at 20 ºC at 3,000 rpm, the rotor was accelerated to 35,000 rpm and radial scans of the sample were performed at 1 min interval. Apparent sedimentation coefficient distributions were calculated using the time-derivative method^24^ with the Origin software provided by the instrument manufacturer. Observed sedimentation coefficient values were converted to the values corresponding to the density and viscosity of water at 20 ºC (s_20,w_) using Sednterp software.

### Electron microscopy

The grids for negative staining electron microscopy were prepared according to previously published protocols.^28^ In brief, 3 μl of SKD3 with 1 mM ATP was applied to glow-discharged 400 mesh formvar carbon film grids with copper supports (Electron Microscopy Sciences, EMS) and stained with a 5 mg/ml uranyl formate solution. For transmission electron microscopy, a total of 45 micrographs were manually acquired using a JOEL JEM-1400 transmission electron microscope with an acceleration voltage of 100 kV. Images were collected at 30,000x magnification with a physical pixel size of 3.897 Å and nominal dosage of 35 e^-^/Å^2^. Image processing was performed using Cryosparc v3.2.0.^29^ The 45 micrographs were CTF-corrected, and 121 particles were manually picked to build an initial 2D set. Using this template, a total of 138,410 unique particles were picked and subjected to 2D classification.

## Acknowledgments

We thank Dr. Sheila Barros for help with DLS experiments and Dr. Sunitha Shiva for help with the enzymatic activity assay. This research was supported by grants from the National Institutes of Health, R01AI141586 (to M.Z.) and from the Johnson Cancer Research Center (to A.Z.).

## Author contributions

M.Z., A.Z., and L.Z., designed the experiments; Z.S., I.T., L.G.S., and M.Z. performed the experiments and analyzed the results; M.Z, A.Z, and L.Z. wrote and edited the manuscript.

## References

1. Neuwald, A. F., Aravind, L., Spouge, J. L., Koonin, E. V. (1999). AAA+: A class of chaperone-like ATPases associated with the assembly, operation, and disassembly of protein complexes. Genome Res. 9, 27–43.

2. Hanson, P. I., Whiteheart, S. W. (2005). AAA+ proteins: have engine, will work. Nat. Rev. Mol. Cell Biol. 6, 519–29.

3. Erives, A. J., Fassler, J. S. (2015). Metabolic and chaperone gene loss marks the origin of animals: evidence for Hsp104 and Hsp78 chaperones sharing mitochondrial enzymes as clients. PLoS ONE 10, e0117192.

4. Cupo, R. R., Shorter, J. (2020). Skd3 (human ClpB) is a potent mitochondrial protein disaggregase that is inactivated by 3-methylglutaconic aciduria-linked mutations. eLife 9, e55279.

5. Seyffer, F., Kummer, E., Oguchi, Y., Winkler, J., Kumar, M., Zahn, R., Sourjik, V., Bukau, B., Mogk, A. (2012). Hsp70 proteins bind Hsp100 regulatory M domains to activate AAA+ disaggregase at aggregate surfaces. Nat. Struct. Mol. Biol. 19, 1347–55.

6. Rosenzweig, R., Moradi, S., Zarrine-Afsar, A., Glover, J. R., Kay, L. E. (2013). Unraveling the mechanism of protein disaggregation through a ClpB-DnaK interaction. Science 339, 1080–3.

7. Thevarajan, I., Zolkiewski, M., Zolkiewska, A. (2020). Human CLPB forms ATP-dependent complexes in the mitochondrial intermembrane space. Int. J. Biochem. Cell Biol. 127, 105841.

8. Saita, S., Nolte, H., Fiedler, K. U., Kashkar, H., Venne, A. S., Zahedi, R. P., Kruger, M., Langer, T. (2017). PARL mediates Smac proteolytic maturation in mitochondria to promote apoptosis. Nat. Cell Biol. 19, 318–328.

9. Wortmann, S. B., Zietkiewicz, S., Kousi, M., Szklarczyk, R., Haack, T. B., Gersting, S. W., Muntau, A. C., Rakovic, A., Renkema, G. H., Rodenburg, R. J., Strom, T. M., Meitinger, T., Rubio-Gozalbo, M. E., Chrusciel, E., Distelmaier, F., Golzio, C., Jansen, J. H., van Karnebeek, C., Lillquist, Y., Lucke, T., Ounap, K., Zordania, R., Yaplito-Lee, J., van Bokhoven, H., Spelbrink, J. N., Vaz, F. M., Pras-Raves, M., Ploski, R., Pronicka, E., Klein, C., Willemsen, M. A., de Brouwer, A. P., Prokisch, H., Katsanis, N., Wevers, R.A. (2015). CLPB mutations cause 3-methylglutaconic aciduria, progressive brain atrophy, intellectual disability, congenital neutropenia, cataracts, movement disorder. Am. J. Hum. Genet. 96, 245–57.

10. Capo-Chichi, J. M., Boissel, S., Brustein, E., Pickles, S., Fallet-Bianco, C., Nassif, C., Patry, L., Dobrzeniecka, S., Liao, M., Labuda, D., Samuels, M. E., Hamdan, F. F., Vande Velde, C., Rouleau, G. A., Drapeau, P., Michaud, J. L. (2015). Disruption of CLPB is associated with congenital microcephaly, severe encephalopathy and 3-methylglutaconic aciduria. J. Med. Genet. 52, 303–11.

11. Kanabus, M., Shahni, R., Saldanha, J. W., Murphy, E., Plagnol, V., Hoff, W. V., Heales, S., Rahman, S. (2015). Bi-allelic CLPB mutations cause cataract, renal cysts, nephrocalcinosis and 3-methylglutaconic aciduria, a novel disorder of mitochondrial protein disaggregation. J. Inherit. Metab. Dis. 38, 211–9.

12. Saunders, C., Smith, L., Wibrand, F., Ravn, K., Bross, P., Thiffault, I., Christensen, M., Atherton, A., Farrow, E., Miller, N., Kingsmore, S. F., Ostergaard, E. (2015). CLPB variants associated with autosomal-recessive mitochondrial disorder with cataract, neutropenia, epilepsy, and methylglutaconic aciduria. Am. J. Hum. Genet. 96, 258–65.

13. Pronicka, E., Ropacka-Lesiak, M., Trubicka, J., Pajdowska, M., Linke, M., Ostergaard, E., Saunders, C., Horsch, S., van Karnebeek, C., Yaplito-Lee, J., Distelmaier, F., Ounap, K., Rahman, S., Castelle, M., Kelleher, J., Baris, S., Iwanicka-Pronicka, K., Steward, C. G., Ciara, E., Wortmann, S. B., Additional individual, c. (2017). A scoring system predicting the clinical course of CLPB defect based on the foetal and neonatal presentation of 31 patients. J. Inherit. Metab. Dis. 40, 853–860.

14. Wortmann, S. B., Zietkiewicz, S., Guerrero-Castillo, S., Feichtinger, R. G., Wagner, M., Russell, J., Ellaway, C., Mroz, D., Wyszkowski, H., Weis, D., Hannibal, I., von Stulpnagel, C., Cabrera-Orefice, A., Lichter-Konecki, U., Gaesser, J., Windreich, R., Myers, K. C., Lorsbach, R., Dale, R. C., Gersting, S., Prada, C. E., Christodoulou, J., Wolf, N. I., Venselaar, H., Mayr, J. A., Wevers, R. A. (2021). Neutropenia and intellectual disability are hallmarks of biallelic and de novo CLPB deficiency. Genet. Med. 23, 1705–1714.

15. Warren, J. T., Cupo, R. R., Wattanasirakul, P., Spencer, D., Locke, A. E., Makaryan, V., Bolyard, A. A., Kelley, M. L., Kingston, N. L., Shorter, J., Bellanne-Chantelot, C., Donadieu, J., Dale, D. C., Link, D. C. (2021). Heterozygous Variants of CLPB are a Cause of Severe Congenital Neutropenia. Blood https://doi.org/10.1182/blood.2021010762.

16. Chen, X., Glytsou, C., Zhou, H., Narang, S., Reyna, D. E., Lopez, A., Sakellaropoulos, T., Gong, Y., Kloetgen, A., Yap, Y. S., Wang, E., Gavathiotis, E., Tsirigos, A., Tibes, R., Aifantis, I. (2019). Targeting Mitochondrial Structure Sensitizes Acute Myeloid Leukemia to Venetoclax Treatment. Cancer Discov 9, 890–909.

17. Glaza, P., Ranaweera, C. B., Shiva, S., Roy, A., Geisbrecht, B. V., Schoenen, F. J., Zolkiewski, M. (2021). Repurposing p97 inhibitors for chemical modulation of the bacterial ClpB-DnaK bichaperone system. J Biol Chem 296, 100079.

18. Lee, S., Sowa, M. E., Watanabe, Y. H., Sigler, P. B., Chiu, W., Yoshida, M., Tsai, F. T. (2003). The structure of ClpB: a molecular chaperone that rescues proteins from an aggregated state. Cell 115, 229–40.

19. Akoev, V., Gogol, E. P., Barnett, M. E., Zolkiewski, M. (2004). Nucleotide-induced switch in oligomerization of the AAA+ ATPase ClpB. Protein Sci. 13, 567–74.

20. Vale, R. D. (2000). AAA proteins. Lords of the ring. J. Cell Biol. 150, F13–9.

21. Lee, S., Choi, J. M., Tsai, F. T. (2007). Visualizing the ATPase cycle in a protein disaggregating machine: structural basis for substrate binding by ClpB. Mol. Cell 25, 261–71.

22. Yokom, A. L., Gates, S. N., Jackrel, M. E., Mack, K. L., Su, M., Shorter, J., Southworth, D. R. (2016). Spiral architecture of the Hsp104 disaggregase reveals the basis for polypeptide translocation. Nat. Struct. Mol. Biol. 23, 830–7.

23. Lopez, K. E., Rizo, A. N., Tse, E., Lin, J., Scull, N. W., Thwin, A. C., Lucius, A. L., Shorter, J., Southworth, D. R. (2020). Conformational plasticity of the ClpAP AAA+ protease couples protein unfolding and proteolysis. Nat. Struct. Mol. Biol. 27, 406–416.

24. Stafford, W. F., 3rd. (1992). Boundary analysis in sedimentation transport experiments: a procedure for obtaining sedimentation coefficient distributions using the time derivative of the concentration profile. Anal. Biochem. 203, 295–301.

25. Jessop, M., Felix, J., Gutsche, I. (2021). AAA+ ATPases: structural insertions under the magnifying glass. Curr. Opin. Struct. Biol. 66, 119–128.

26. Gates, S. N., Martin, A. (2020). Stairway to translocation: AAA+ motor structures reveal the mechanisms of ATP-dependent substrate translocation. Protein Sci 29, 407–419.

27. Cupo, R. R., Shorter, J. (2020). Expression and Purification of Recombinant Skd3 (Human ClpB) Protein and Tobacco Etch Virus (TEV) Protease from Escherichia coli. Bio Protoc 10, e3858.

28. Peisley, A., Skiniotis, G. (2015). 2D Projection Analysis of GPCR Complexes by Negative Stain Electron Microscopy. Methods Mol. Biol. 1335, 29–38.

29. Punjani, A., Rubinstein, J. L., Fleet, D. J., Brubaker, M. A. (2017). cryoSPARC: algorithms for rapid unsupervised cryo-EM structure determination. Nat. Methods 14, 290–296.

